# 5-Fluorouracil Treatment Represses Pseudouridine-Containing Small RNA Export into Extracellular Vesicles

**DOI:** 10.1101/2024.01.15.575751

**Authors:** Sherman Qu, Hannah Nelson, Xiao Liu, Elizabeth Semler, Danielle L. Michell, Clark Massick, Jeffrey L. Franklin, John Karijolich, Alissa M. Weaver, Robert J. Coffey, Qi Liu, Kasey C. Vickers, James G. Patton

**Author notes:** Corresponding authors: James G Patton Department of Biological Sciences, Vanderbilt University Kasey C. Vickers Department of Medicine, Vanderbilt University Medical Center. These authors contributed equally to this manuscript.

## Abstract

5-fluorouracil (5-FU) has been used for chemotherapy for colorectal and other cancers for over 50 years. The prevailing view of its mechanism of action is inhibition of thymidine synthase leading to defects in DNA replication and repair. However, 5-FU is also incorporated into RNA causing toxicity due to defects in RNA metabolism, inhibition of pseudouridine modification, and altered ribosome function. Here, we examine the impact of 5-FU on the expression and export of small RNAs (sRNAs) into small extracellular vesicles (sEVs). Moreover, we assess the role of 5-FU in regulation of post-transcriptional sRNA modifications (PTxM) using mass spectrometry approaches. EVs are secreted by all cells and contain a variety of proteins and RNAs that can function in cell-cell communication. PTxMs on cellular and extracellular sRNAs provide yet another layer of gene regulation. We found that treatment of the colorectal cancer (CRC) cell line DLD-1 with 5-FU led to surprising differential export of miRNA snRNA, and snoRNA transcripts. Strikingly, 5-FU treatment significantly decreased the levels of pseudouridine on both cellular and secreted EV sRNAs. In contrast, 5-FU exposure led to increased levels of cellular sRNAs containing a variety of methyl-modified bases. Our results suggest that 5-FU exposure leads to altered expression, base modifications, and mislocalization of EV base-modified sRNAs.

## Introduction

Extracellular vesicles (EVs) are membrane-bound particles that vary in size, cargo composition, biogenesis pathways, and delivery mechanisms (Couch et al. 2021; Dixson et al. 2023). The demonstration that EVs released from B cells could functionally activate T cells accelerated interest in the EV field, which was further increased by findings that EVs contain protein and RNA cargo that can be transferred between donor and recipient cells (Raposo et al. 1996; Thery et al. 2002; Ratajczak et al. 2006; Valadi et al. 2007; Skog et al. 2008). It is now appreciated that all cells release EVs and that EVs can constitute a unique form of cell-cell communication, from near-neighbor interactions to communication between distant cells and organs (Maas et al. 2017; van Niel et al. 2018; Wortzel et al. 2019; Couch et al. 2021; Buzas 2023; Dixson et al. 2023).

Membrane trafficking studies using reticulocytes showed by electron microscopy that intraluminal vesicles contained within multivesicular bodies (MVBs or late endosomes) are released from cells and the term exosomes was coined in the 1980s to describe such particles (Harding et al. 1983; Pan et al. 1985). Heterogeneous populations of EVs are released from cells using multiple biogenesis pathways with current focus to more precisely name the different classes of vesicles based on size and biogenesis pathway (Colombo et al. 2014; Abels and Breakefield 2016; Thery et al. 2018; Jeppesen et al. 2023). As it is difficult to identify a biogenesis pathway from standard EV preparations, we will use the term small EVs (sEV) to describe our vesicle preparations. Non-vesicular particles and lipoproteins are also released from cells that can transfer protein and RNA cargo between cells (Vickers et al. 2011; Zhang et al. 2018; Jeppesen et al. 2019; Zhang et al. 2019; Zhang et al. 2021; Allen et al. 2022). Here, we investigate the role of post-transcriptional modifications (PTxM) on sRNA selection and/or export in response to 5-FU treatments of CRC cells.

Protein, RNA and lipid cargo associated with EVs varies in a cell-context dependent manner (Tkach and Thery 2016; Maas et al. 2017; Dixson et al. 2023). Almost all known classes of coding and noncoding RNA (mRNA, rRNA, tRNA, snRNA, snoRNA, Y RNAs, ncRNAs) have been detected in EVs (Nolte-’t Hoen et al. 2012; Cha et al. 2015; Fritz et al. 2016; Turchinovich et al. 2019; Veziroglu and Mias 2020; Dellar et al. 2022). The majority of RNAs within EVs correspond to small RNAs and fragments of larger transcripts (<200nt) (Mateescu et al. 2017; Turchinovich et al. 2019). Among small RNAs, microRNAs (miRNAs) constitute one of the best characterized RNA components in EVs because they exist as full length RNAs within EVs and because their transfer between cells can be assayed using standard reporter assays (Cha et al. 2015). Numerous studies have proposed that miRNA export into EVs is regulated (Villarroya-Beltri et al. 2013; Squadrito et al. 2014; Cha et al. 2015; McKenzie et al. 2016; Santangelo et al. 2016; Shurtleff et al. 2016; Shurtleff et al. 2017). Most regulatory hypotheses have proposed that specific sequence motifs promote selective export by candidate RNA binding proteins, but no universal targeting sequence has thus far been identified (Bolukbasi et al. 2012; Villarroya-Beltri et al. 2013; Santangelo et al. 2016; Shurtleff et al. 2016; Temoche-Diaz et al. 2019; Garcia-Martin et al. 2022).

Over 200 hundred post-transcriptional base modifications have been identified along with many writers, readers, and erasers of such modifications (Roundtree et al. 2017; Schaefer et al. 2017; Zaccara et al. 2019). We posit that PTxMs on sRNAs serve as export signals for EV sRNAs (Abner et al. 2021). Recently, we found that knockdown of Mettl3, a writer of N6-methyladenosine (m^6^A) modification, reduced EV levels of miRNAs that contain consensus sequences for m^6^A (Abner et al. 2021). Here we identify a large number of PTxMs on sRNAs with pseudouridine being the most abundant. To determine the effect of altered pseudouridine modification on sEV sRNA export, we treated CRC cells with 5-fluorouracil (5-FU), an inhibitor of pseudouridine modification (Samuelsson 1991; Gu et al. 1999; Zhao and Yu 2007; Karijolich et al. 2010; Liang et al. 2022). 5-FU has been used in cancer treatment for >50 years with the presumed mechanism of action being inhibition of thymidine synthesis to block DNA replication (Longley et al. 2003). However, 5-FU is also incorporated into RNA and recent work has shown that the effects of 5-FU are more complex, with effects on a variety of RNAs, some of which lead to altered ribosome function and translational recoding (Mojardin et al. 2013; Ge et al. 2017; Chalabi-Dchar et al. 2021; Therizols et al. 2022). One possible mechanism to explain RNA-mediated effects after incorporation of 5-FU is through inhibition of one of more of the 13 pseudouridine synthases (Jin et al. 2022). Remarkably, we find that 5-FU treatment dramatically decreased the levels of pseudouridine in both cells and sEVs and led to decreased export of miRNAs and sRNA fragments derived from spliceosomal RNAs. Opposite effects were observed for sRNAs derived from snoRNAs. Together, our results show that 5-FU treatment leads to changes in cellular gene expression, massive changes in PTxM status on cellular sRNAs, and loss of selective EV export of miRNAs, and snRNA- and snRNA-derived fragments.

## Materials and Methods

### Cell Culture

DLD-1 cells (Shirasawa et al. 1993) were validated as heterozygous KRAS mutant (G13D) cells by PCR amplification of genomic DNA and DNA sequencing. Cells were seeded at 6.5x 10^6^ in T175 flasks and cultured in DMEM medium with 10% fetal bovine serum, 2 mM L-glutamine, 1x MEM non-essential amino acids, 100 U/mL of penicillin, and 100 U/mL of streptomycin. When the cells reached 70% confluence, they were washed three times with phosphate-buffered saline (PBS) and cultured in a serum-free medium with or without 10 μM 5-FU for 48 hrs.

### Extracellular Vesicle Isolation

Media from at least three T175 flasks were pooled for extracellular vesicle (EV) isolation as in described (Hinger et al. 2020; Abner et al. 2021). Briefly, conditioned media was centrifuged at 1000xg for 10 minutes at room temperature. Supernatants were transferred to a new centrifuge tube and spun at 2,000xg for 10 minutes at 4°C after which the supernatant was again transferred to a new centrifuge tube and spun at 10,000xg for 30 minutes at 4°C. The supernatants were collected, transferred to clean ultracentrifuge tubes and spun at 167,000xg for 17 hours at 4°C. The resulting pellets were washed twice by resuspension 1 ml sterile Dulbecco’s PBS and pelleting at 167,000xg for 70 minutes at 4°C. After the final wash, pellets were resuspended in 50μL of sterile 1X DPBS.

### RNA Isolation

Cells were collected by scraping, washed in PBS, and resuspended in PBS prior to isolation of total RNA using the Quick-RNA Miniprep kit (Zymo Research). Total RNA from pelleted EVs in PBS was isolated using the same kit.

### Small RNA sequencing (sRNA-seq)

sRNA libraries were generated using the NEXTFlex Small RNA Library Preparation Kits v3 (Perkin) with the following modifications: (1) 3′- and 5′-adaptors were diluted 1:8, (2) 3′-adaptor ligations were performed overnight in multiple steps – 25°C for 2 h, 20°C for 4 h and 16°C overnight, (3) following cDNA synthesis and before barcoding PCR, step F protocol was followed, and (4) PCR amplification was 20 cycles. Following PCR amplification, individual libraries were size-selected (136–200 bp product) using Pippin Prep (Sage Sciences). Size-selected libraries were quantified using Qubit Fluorometer. Libraries were checked for quality and sequenced using Illumina short-read technology. Libraries were pooled and paired-end sequencing (PE- 150) of the pool (equimolar multiplexed libraries) was performed on the NovaSeq6000 platform by the VANTAGE core (Vanderbilt University). Demultiplexing and bioinformatic analyses was performed using the TIGER pipeline (Allen et al. 2018). Briefly, Cutadapt (v1.16) was used to trim 3′ adaptors and all reads with < 16 nucleotides (nts) were removed (Martin 2011). Quality control on both raw reads and adaptor-trimmed reads was performed using FastQC (v0.11.9)(www.bioinformatics.babraham.ac.uk/projects/fastqc). The adaptor-trimmed reads were mapped to the hg19 genome, with additional rRNA and tRNA reference sequences, by Bowtie1 (v1.1.2) allowing only one mismatch (Langmead and Salzberg 2012).

### Total RNA Sequencing

Bulk RNA sequencing libraries were prepared using Universal RNAseq kits (Tecan), as per manufacturer’s instructions. Libraries were cleaned (Zymo), checked for quality using the Agilent bioanalyzer, quantified (Qubit), and pooled based on equimolar concentrations of libraries. Pooled libraries were sequenced using Illumina short-read technology based on PE150 on the NovaSeq6000 (Illumina). After sequencing, samples (libraries) were demultiplexed and analyzed. Briefly, reads were trimmed to remove adapter sequences using Cutadapt v2.10)(Martin 2011) and aligned to the Gencode GRCh38.p13 genome using STAR (v2.7.8a)(Dobin et al. 2013). Gencode v38 gene annotations were provided to STAR to improve the accuracy of mapping. Quality control on both raw reads and adaptor-trimmed reads was performed using FastQC (v0.11.9). FeatureCounts (v2.0.2)(Liao et al. 2014) was used to count the number of mapped reads to each gene. Heatmap3 (Zhao et al. 2014) was used for cluster analysis and visualization. Significant differential expressed genes with absolute fold change > 2 or < 0.5, and adjusted p value (padj) < 0.05 were determined using DESeq2 (v1.30.1)(Love et al. 2014). Genome Ontology and KEGG pathway over- representation analyses were performed on differentially expressed genes using the WebGestaltR package (NULL)(Wang et al. 2017). Gene set enrichment analysis (GSEA) was performed using GSEA package (v4.3.2)(Subramanian et al. 2005) on database v2022.1.Hs.

### Mass Spectrometry

Liquid chromatography tandem mass spectrometry (LC-MS/MS) was used to quantify PTxMs based on standard curves. Main nucleosides--adenosine (A), cytidine (C), guanosine (G) and uridine (U)--were obtained from Sigma-Aldrich. 1- methyladenosine (m^1^A), 2-methyladenosine (m^2^A), 6-methyladenosine (m^6^A), 8- methyladenosine (m^8^A), 3-methylcytidine (m^3^C), pseudouridine, 5-methylcytidine (m^5^C), 5-methyluridine (m^5^U), 1-methylguanosine (m^1^G), 2-methylguanosine (m^2^G), 2- dimethylguanosine (m_2_ ^2^G), 7-methylguanosine (m^7^G) were purchased from Carbosynth. [^15^N]_5_-2-deoxyadenosine was obtained from Cambridge Isotope Laboratories. Cellular and EV RNAs were enzymatically digested to single nucleosides under neutral conditions using the Nucleoside Digestion Mix (BioLabs). RNA nucleosides were combined with 1 nM of [^15^N]-dA internal standard immediately prior to analysis. Digested nucleosides were separated using a Hypersil GOLD aQ C18 column (100-mm length × 2.1-mm inner diameter, pore size 175 Å, particle size 1.9 µm; (ThermoFisher) at a flow rate of 0.4 mL/min using 0.1% (v/v) formic acid in water (solvent A) and acetonitrile with 0.1% (v/v) formic acid (solvent B). The gradient profile applied to each sample was as follows: 0–6 min, 0% B; 6–7.5 min, 1% B; 7.5-9.5 min, 6% B; 9.5-10.5 min, 10.5-12 min, 50% B; 12-14 min, 75% B; 14-16 min, 75% B. The column was equilibrated prior to subsequent injections (10 µL) and maintained at 40°C. MS analysis was performed on a Shimadzu Nexera system in-line with a QTRAP 6500 with an electrospray ionization source (Applied Biosystems). Multiple reaction monitoring (MRM) in positive ion mode was used to survey known modifications. Data acquisition and sample processing were performed using AB SCIEX Analyst Software 1.6.2 (Applied Biosystems). Nucleosides were quantified by converting peak area under the curve to nmol using standard curves of candidate nucleosides. To compare modification levels across conditions, modified nucleoside values were normalized to total nucleosides analyzed.

### Statistics

To compare between 2 groups, two-tailed unpaired t-tests were used with p<0.05 considered significant. The Wilcox-rank sum test was used to compare U percentages between miRNAs with significant interaction effect and those without.

## Results

To determine the effects of 5-FU treatment on gene expression, we exposed the patient-derived CRC cell line DLD-1 (Shirasawa et al. 1993) to 10μM 5-FU for 48 hours in serum-free media to avoid EV contamination. Total RNA was purified from both cell lysates and secreted EVs and RNA sequencing was performed on 5-FU treated cells and controls (**Table S1**).

### Total RNAseq

We first analyzed total RNAseq data from DLD-1 cells in the presence (FiveFU) or absence (SF) of 5-FU exposure. From the raw sequencing data, 5-FU treatment caused significant (adjusted p<0.05) differential (absolute fold change >2.0) expression of 807 transcripts with 365 up- and 442 down-regulated RNAs when comparing treated to untreated cells (**Fig. 1A, B**)(**Table S1**). When expression threshold levels were applied to restrict analysis to mean read values of at least 40, there were 345 up- regulated and 324 down-regulate genes.

**Figure 1.**
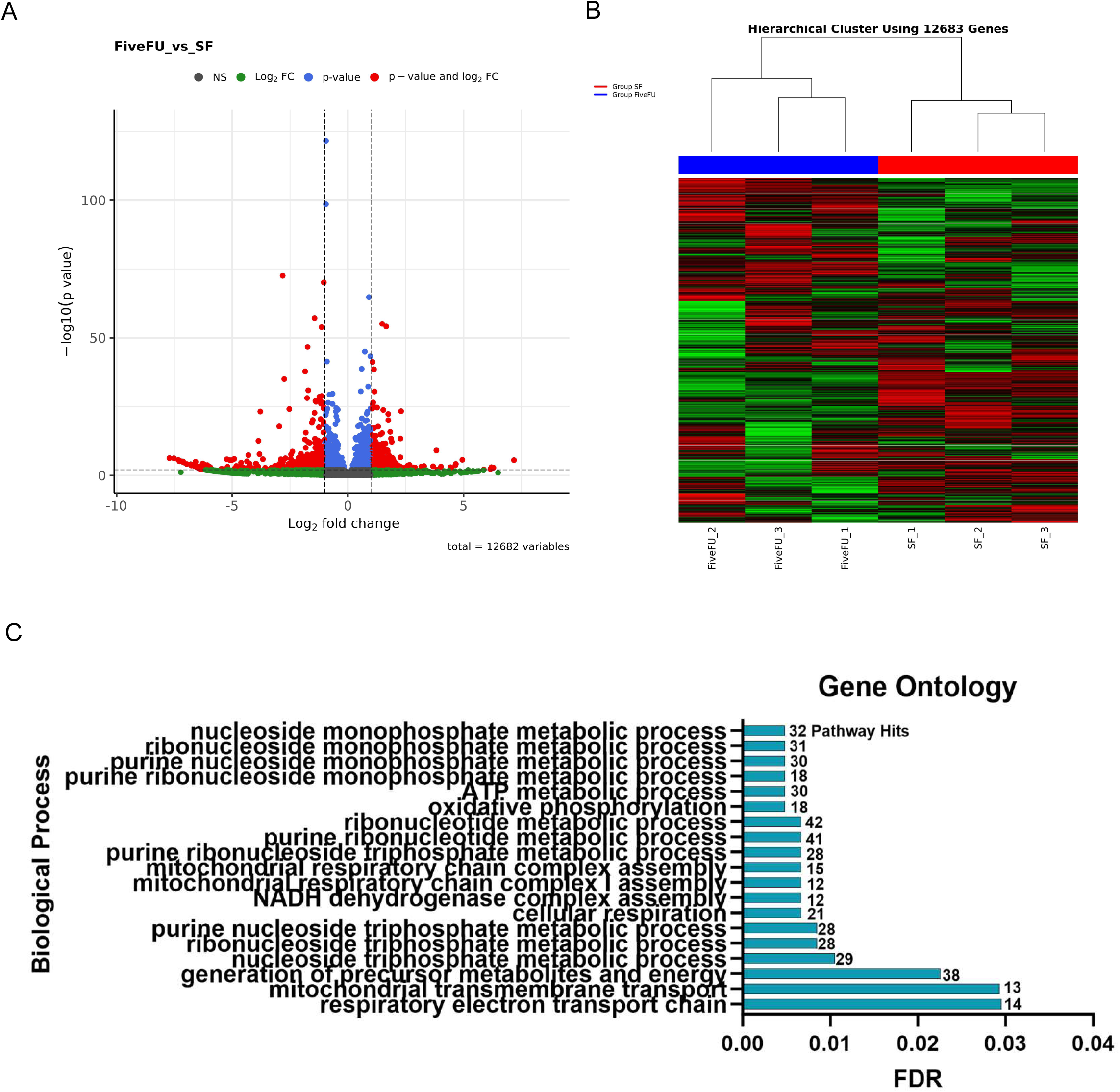
Differential gene expression in DLD-1 cells after exposure to 5-FU. DLD-1 cells were exposed (FiveFU) or not (SF) to 10uM 5-FU for 48hr in the absence of serum after which RNA was isolated and RNAseq was performed on transcripts >200nt. **A**. Volcano plot showing up- and down-regulated genes after treatment with 5-FU. The identity of individual data points are in Table S1. Red dots represent transcripts with at least 2-fold changes (up and down) in expression and adjusted p values < 0.05. Blue dots represent transcripts with adjusted p values <0.05 but whose expression changes were less than 2-fold. Green dots represent transcripts with 2-fold changes in gene expression but adjusted p values >0.05. Black dots represent transcripts whose expression did not significantly change after 5-FU treatment. **B**. Hierarchical clustering analysis was performed on triplicate RNAseq samples prepared from DLD-1 cells treated (blue, FiveFU) or not (red, SF) with 5-FU. **C**. Gene ontology analysis was performed using the WebGestaltR package (Wang et al. 2017).

To identify specific pathways and processes that were affected by 5-FU treatment, Gene Ontology analysis showed significant enrichment of nucleoside metabolism pathways (**Fig. 1C).** Also, several RNA metabolism pathway genes (GO:0016070) were found to be significantly increased (FDR=1.5x10^-5^) by 5-FU treatment with 136 RNA metabolism-associated genes significantly increased at the mRNA level (**Tables S1, S2)**. This includes several PTxM enzymes: tRNA methyltransferase 61 (*TRM61*, FC=2.17, p=0.004), Dihydrouridine Synthase 1 Like 1 (*DUSL1*, FC=2.25, p=0.0000073), c16orf42 Ribosome Maturation Factor (*TSR3M,* FC=2.17, p=0.011), and Pseudouridine Synthase 7 (*PUS7*, FC=2.15, p=0.0023). These observations are consistent with previous reports of thymidine synthase inhibition and DNA replication defects by 5-FU, but also with more recent findings that RNA-based defects also underlie the mechanism of action of 5-FU (Mojardin et al. 2013; Ge et al. 2017; Chalabi-Dchar et al. 2021; Liang et al. 2022; Therizols et al. 2022).

### Small RNAseq

To investigate the impact of 5-FU on sRNA expression and export in sEVs, high-throughput sRNA-seq was completed on RNA isolated from donor DLD-1 cells and secreted sEVs in control and 5-FU treated conditions (**Table S3).** Cellular and EV sRNAs were quantified using short-read Illumina sequencing with degenerate base adapters for library generation. The data met quality control metrics as analyzed using the TIGER sRNA pipeline (Allen et al. 2018)(**Fig. 2A, Fig. S1A,B**). In cells, host sRNA counts per sample averaged 17.5 million read per EV samples (**Fig. 2A**). The most abundant class of host sRNAs detected in cells were miRNAs followed by sRNAs derived from parent rRNAs (rDR) (**Fig. 2A**). In contrast, the most abundant host sRNA classes in secreted EVs were tRNA-derived sRNAs (tDRs) and Y RNA-derived sRNAs (yDR) (**Fig. 2A**). Hierarchical clustering and correlation analyses were used to show the lack of association between cellular and EV miRNA profiles (**Fig. 2B**).

**Figure 2.**
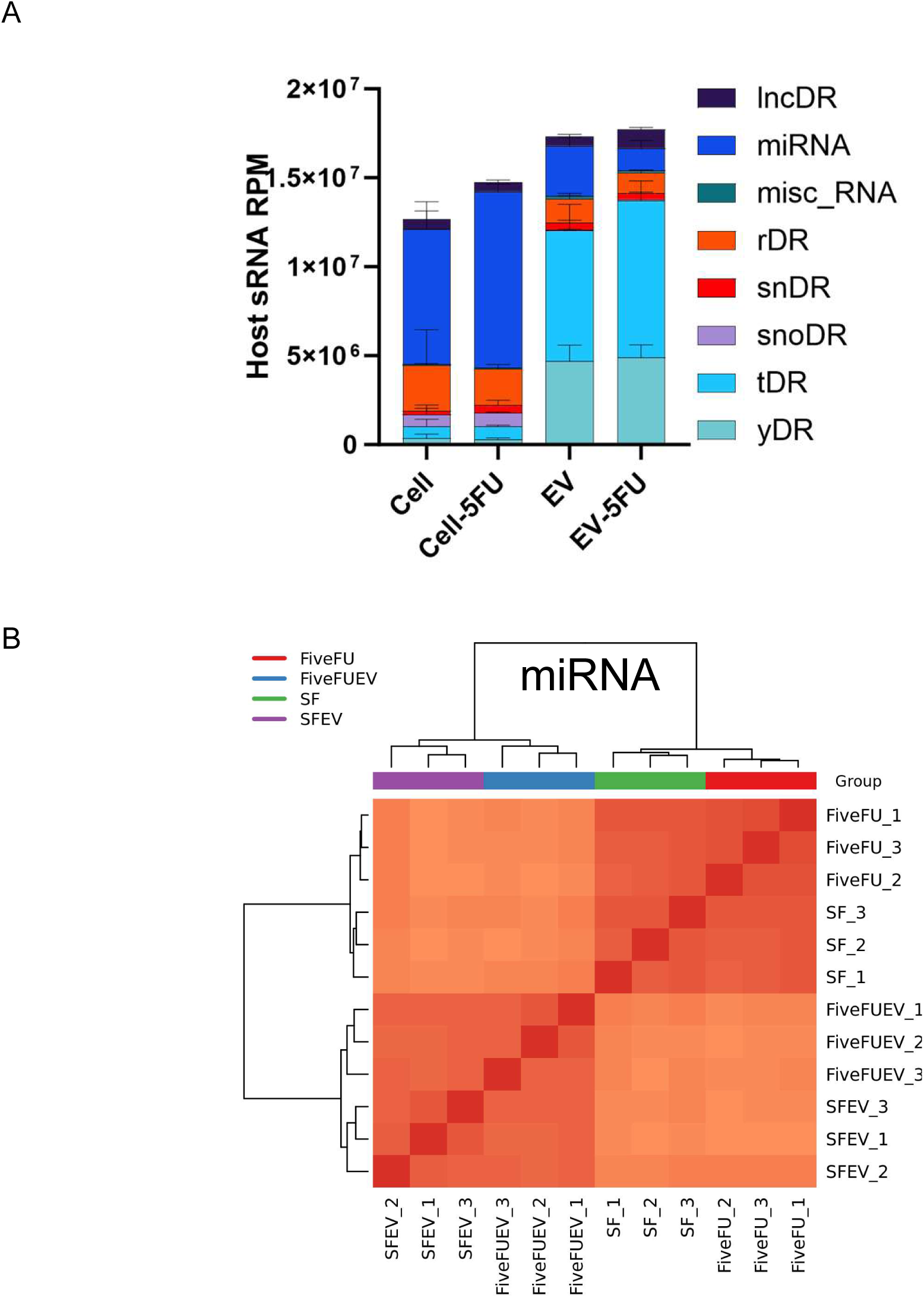
Small RNAseq of DLD-1 cells and extracellular vesicles treated with 5-FU. Small RNAseq was performed on RNA (<200nt) isolated from DLD-1 cells and EVs treated or not with 5-FU. **A**. Total number of small RNA (sRNA) reads and identity of reads from the indicated RNA subclasses after RNAseq on cellular and EV RNA isolated in the presence or absence of 5-FU. RNA subclasses include long noncoding RNA-derived sRNAs (lncDRs), miRNAs, miscellaneous sRNAs, rRNA-derived sRNAs (rDRs), spliceosomal snRNA-derived sRNAs (snDRs), snoRNA- derived sRNAs (snoDRs), tRNA-derived sRNAs (tDRs) and Y RNA-derived fragments (yDRs). **B**. Clustering of miRNA expression patterns in cells and EVs after treatment with 5-FU. miRNA expression profiles comparing cells and EVs treated (FiveFU) or not (SF) with 5-FU across triplicate RNAseq samples for each condition.

### Effects of 5-FU on tDR and rDR Expression and EV Export

Non-coding sRNAs harbor many PTxMs, either directly modified or indirectly inherited from their parent transcripts. The functional roles of PTxMs on sRNAs are only beginning to emerge; however, this new level of gene regulation holds great potential for cellular sRNA selection and trafficking for degradation or export. Due to inhibition of pseudouridine synthases by 5-FU (Karijolich et al. 2010), we expected differential cellular and/or EV expression of rDR and tDR sRNAs. Pseudouridine modification of tRNA was one of the first RNA modifications to be characterized, with ∼17 sites now identified (Borchardt et al. 2020). When we examined tDRs in cells treated or not with 5-FU, we found very few differentially expressed tDRs with only 6 up- regulated and no down-regulated tDRs (padj<0.05 and fold-change >1.5 or <0.67)(**Fig. S2, Table S3**). When we examined tDRs in EVs either treated or not with 5-FU, we detected 8 up- and 14 down-regulated tDRs (padj<0.05 and fold-change >1.5 or <0.67)(**Fig. S2, Table S3**). Pseudouridine modification is also prevalent in rRNA with modification sites mapped in E. coli, yeast and human rRNA (Taoka et al. 2018; Borchardt et al. 2020). For rDRs in cells treated with 5-FU, we failed to identify up- regulation of rDRs and only found 2 rDRs to be down-regulated in cells (**Figure S2, Table S3**). For rDRs from EVs purified from cells either treated or not with 5-FU, we failed to detect up-regulated rDRs and only found 4 up-regulated rDRs in EVs (**Figure S2, Table S3**).

### 5-FU inhibits miRNA and snDR export and promotes snoDR export to EVs

In addition to the modest effects of 5-FU on tDR and rDR, we also observed minor effects on yDRs (**Fig. S2**). In contrast, we detected significant changes in expression and/or EV localization for miRNAs, spliceosomal snRNAs (snDRs), and small nucleolar RNA (snoDRs) (**Fig. 3; Table S3**). At the cellular level, distinct subsets of miRNAs were both up- and down-regulated (**Figure 3A**), including 89 up-regulated miRNAs and 42 down-regulated miRNAs after 5-FU treatment (**Table S3)**. This shows that 5-FU treatment affects expression of miRNAs in DLD-1 cells, but with a mix of miRNAs that either up- or down-regulated. In contrast, we observed significant down- regulation of miRNA reads in EVs in the presence of 5-FU (**Figure 3A**), with 254 down- regulated miRNAs (**Table S3**). Combining the cellular and EV data, treatment with 5- FU caused a massive reduction of miRNA reads in EVs, suggesting an export defect.

**Figure 3.**
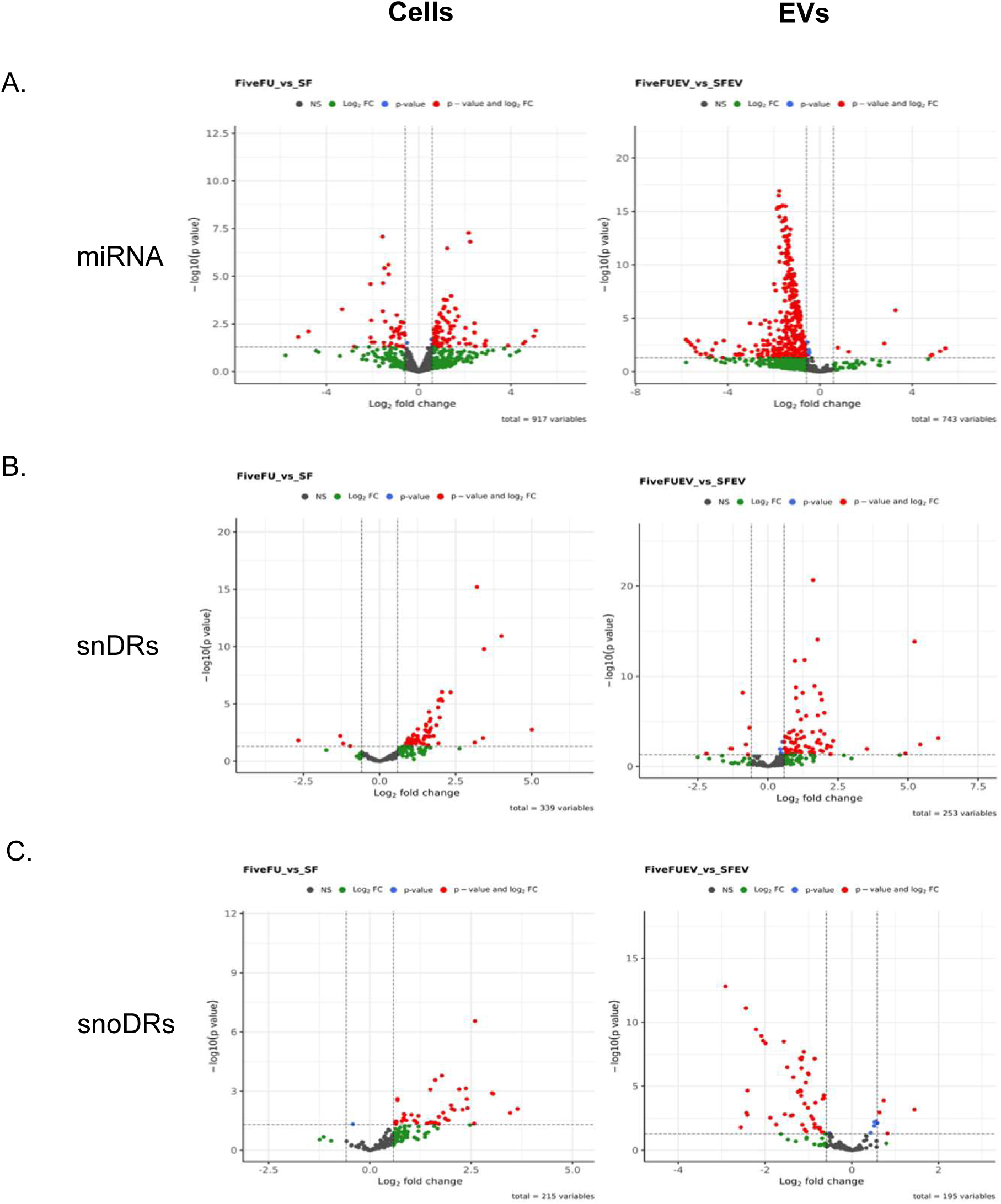
Differentially expressed sRNAs comparing DLD-1 cells and EVs grown in the presence or absence of 5-FU. Volcano plots showing up- and down-regulated miRNAs (**A**), snDRs (**B**) and snoDRs (**C**) in cells and EVs treated (FiveFU) or not (SF) with 5-FU. Individual data points represent transcripts with fold-changes and significance as in Fig. 1.

To determine if any of the observed decreases in DLD-1 EVs could be linked to cellular miRNA changes and sorting defects, interaction effects between the datasets were calculated. We found that many of the significantly decreased EV miRNAs were linked to cellular miRNAs changes (**Table S4**). These results agree with the concept that miRNA sorting and/or selection for export are potentially mediated by PTxMs, which may be regulated by 5-FU therapy.

For snRNAs, we found that many snDRs were up-regulated in cells after treatment with 5-FU; however, despite that increase in expression, there were many down-regulated snDRs in EVs, similar to miRNAs (**Fig. 3B**). For example, 34 snDRs were up-regulated in cells with no detectable down-regulated transcripts in cells treated with 5-FU (**Fig. 3B, Table S3**). In contrast, we observed the opposite effect in EVs with 51 down- and 4 up-regulated snDRs after treatment with 5-FU (**Fig. 3B, Table S3**). This mirrors the miRNA data, with down-regulation of reads in EVs suggesting an export defect in the presence of 5-FU.

For snoRNAs, several snoDRs were up-regulated in cells after treatment with 5- FU, but instead of decreased reads in EVs, as noted for snRNAs and miRNAs, we observed the opposite effect with up-regulation of snoDRs in EVs (**Fig. 3C**). For example, we detected up-regulation of 63 snoDRs and down-regulation of 4 snoDRs in treated cells (**Fig. 3C, Table S3**). In EVs, there were 74 up-regulated and 7 down- regulated snoDRs in EVs (**Fig. 3C, Table S3**).

### Export Defects and Uridine Content

We next decided to examine RNA export and uridine content focusing on the dramatic changes in miRNA export into EVs in the presence of 5-FU. For this, we re-examined differential miRNA expression after 5-FU treatment to identify miRNAs that show contrasting coordinate regulation between cells and EVs meaning down- regulation in cells and up-regulation in EVs or up-regulation in cells and down-regulation in EVs. If we set the fold-changes to >1.8 or <0.5, there were no miRNAs that were both up-regulated in EVs and down-regulated in cells. In contrast, there were 6 miRNAs that were coordinately up-regulated in cells and down-regulated in EVs **(Fig. 4A**). Interestingly, when we examined the base composition of the coordinately regulated miRNAs, all 6 miRNAs contained at least 7 uridine residues across the short mature miRNA sequence and two of these also contain a uridine-rich precursor loop. These data suggest that pseudouridine modification could be important for miRNA export since incorporation of 5-FU at one or more of these positions would inhibit modification. Consistent with this, we performed a Wilcoxon Rank Sum test to compare the percentage of U residues when comparing miRNA populations that were retained in cells or exported to EVs in the presence of 5-FU. We found a statistically significant increase in the percentage of uridines in retained miRNAs (**Fig. 4B**). This analysis supports the idea that pseudouridine modification may be important for RNA export.

**Figure 4.**
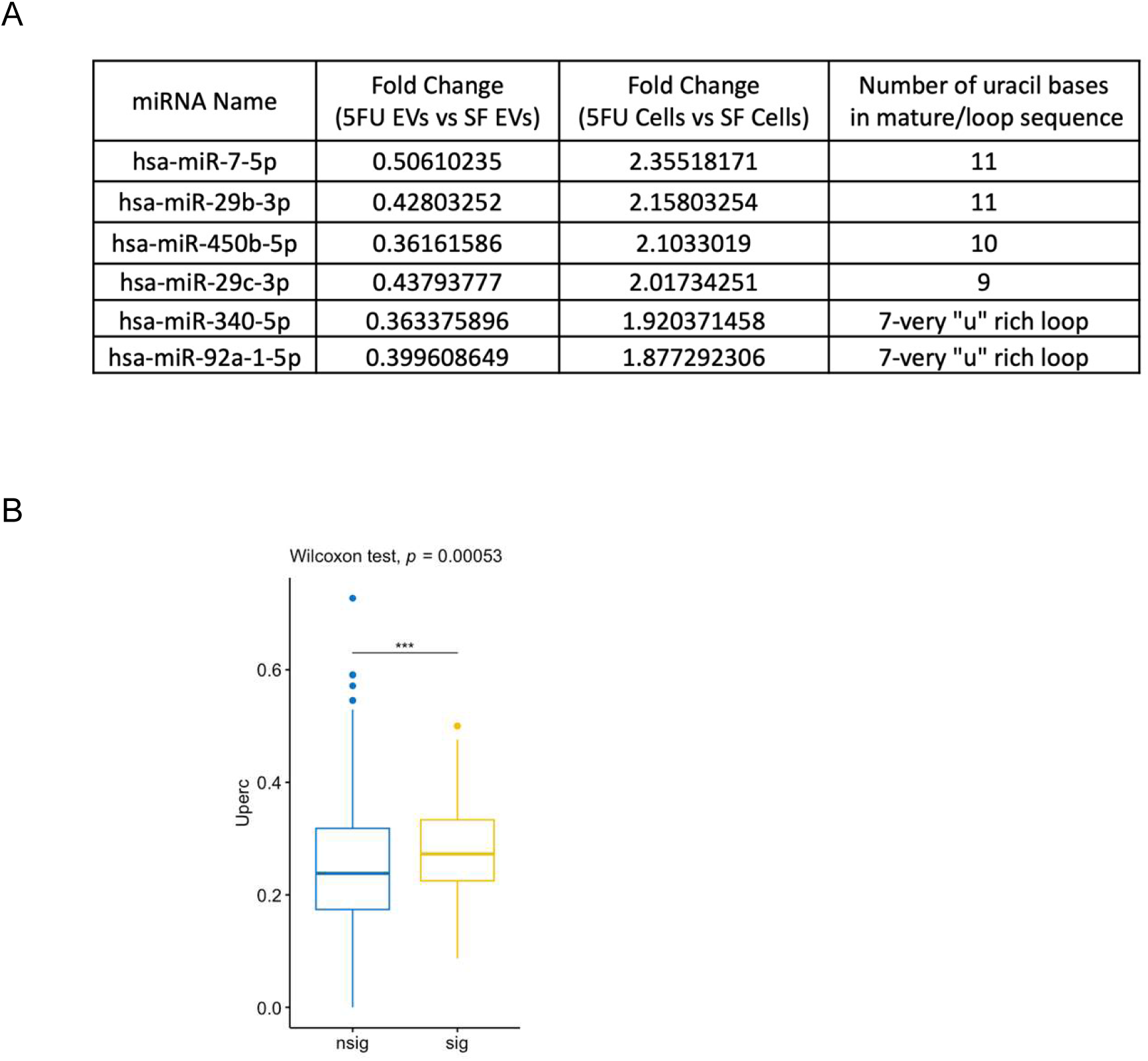
Increased uridine content in cellular-retained miRNAs EV after 5-FU treatment. **A.** miRNAs that display coordinate up-regulation in DLD-1 cells and down-regulation in EVs after treatment with 5-FU were identified. The top miRNAs that show coordinate expression changes after 5-FU are shown along with the number of uridine residues in each mature miRNA. miR-340 and miR-92a also contain uridine rich precursor loop sequences. **B**. Overall enrichment of uridine residues in miRNAs retained in cells after 5-FU treatment. Significant cellular-retained miRNAs in EVs after 5-FU treatment were identified based on criteria comprising a fold change greater than 2 and a false discovery rate (FDR) less than 0.01. The proportion of U content in these significantly retained miRNAs was then compared to that of non-significant ones using the Wilcoxon rank sum test. A significant increase in uridine content was detected in cellular-retained RNAs.

### 5-FU treatment reduces pseudouridine levels in cells and secreted EVs

To determine if 5-FU treatment alters the levels of pseudouridine and other PTxMs, LC-MS/MS approaches were used to quantify candidate PTxM levels in total RNA from cells and secreted sEVs. This allowed quantitation of the levels of the 4 main nucleosides and 14 PTxMs based on internal and external standards. In DLD-1 cells, 5- FU treatments were found to significantly increase uridine (U) levels in cells, but failed to alter the other 3 nucleosides in secreted EVs, as normalized per total nucleoside levels (**Figs. 5A,B**). Importantly, pseudouridine detection significantly decreased in both DLD-1 cells and secreted sEVs with 5-FU treatment (**Fig. 5C**). Conversely, dihydrouridine (DHU) levels were found to be significantly increased in 5-FU treated DLD-1 cells with no observable changes in EVs (**Fig. 5D**). Also, 3-methyluridine (m^3^U) and 5-methyluridine (m^5^U) levels were not altered in 5-FU-treated cells or secreted EVs (**Fig. S3A,B**). The cellular levels of 5-methylcytosine (m^5^C) and 3-methylcytosine (m^3^C) were found to be significantly increased in 5-FU treated cells without changes in sEVs (**Fig. 5E,F**). N1-methyladenosine (m^1^A) levels were also significantly increased in 5-FU treated cells, however, N6-methyladenosine (m^6^A) levels were not affected by 5-FU treatment (**Fig. 5G, Fig. S3C**). In cells, 1-methylinosine (m^1^I) levels were significantly increased with 5-FU treatment, but m^1^I levels were undetectable in secreted sEVs (**Fig. 5H**). Inosine (I) levels were detected in both cells and secreted EVs, however, the levels of I were not affected by 5-FU treatment (**Figure S3D**). For guanosine (G), 1- methylguanosine (m^1^G), 7-methylguanosine (m^7^G), 2-methylguanosine (m^2^G), N2, N2- dimethylguanosine (m^2^ G), and N2, 7-dimethylguanosine (m^2^ G) levels were all significantly increased after treatment with 5-FU (**Fig. I-L, Fig. S3E**). With the exception of m^2^,_7_G, these guanosine PTxMs were also detected in sEVs, however, their levels were not altered by 5-FU (**Fig. 4I-L, Fig. S3E**). These LC-MS/MS data suggest that while post-transcriptional methylation of nucleosides was significantly increased in cells treated with 5-FU, pseudouridine levels were significantly decreased in both treated cells and secreted EVs. As pseudouridine was the only PTxM observed to be altered in EVs with 5-FU treatments, these results support pseudouridine as a factor in sRNA selection and/or export to EVs.

**Figure 5.**
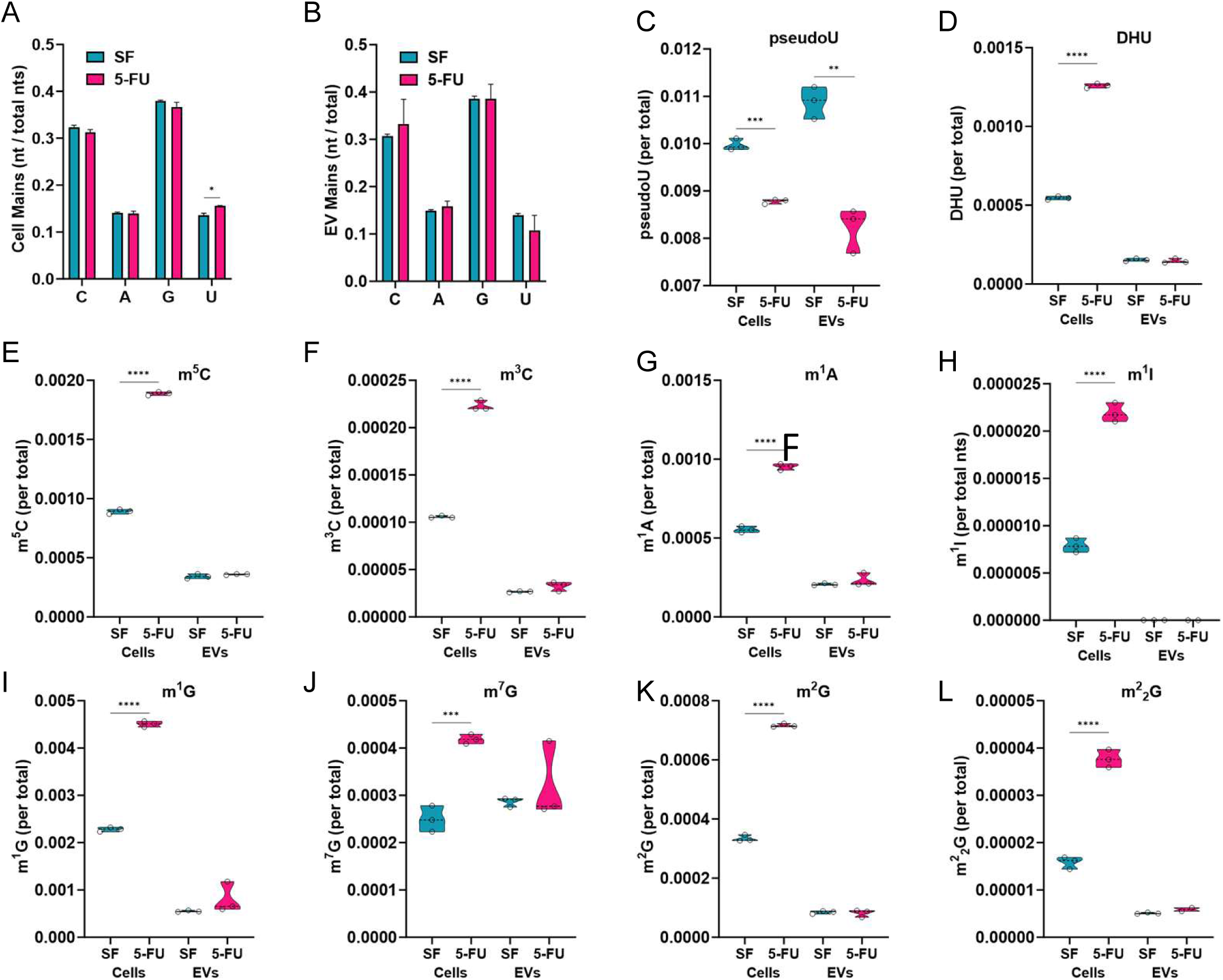
Reduced pseudouridine modification in cells and EVs after 5-FU treatment. LC- MS/MS approaches were used to quantify RNA base modifications in RNA from cells and secreted sEVs. **A.** Nucleoside content in 5-FU treated DLD-1 cells showed a significant increase in uridine content when compared to untreated cells (SF). No changes were observed for the other RNA nucleosides. **B.** No changes in nucleoside levels were detected in EVs purified from cells treated or not with 5-FU. C. Pseudouridine levels were significantly decreased in both cells and EVs after 5-FU treatment. D-L. Quantitation of the effect of the indicated base modifications in cells and EVs treated or not with 5-FU. Base modifications include dihydrouridine (DHU), methylated cytosine (m^5^C and m^3^C), methylated adenosine (m^1^A), methylated inosine (m^1^I), and methylated guanosine (m^1^G, m^7^G, m^2^G) and demethylated guanosine (m^2^_2_G). To compare modification levels across conditions, modified nucleoside values were normalized to total nucleosides.

## Discussion

Here, we report the first analysis of RNA profiles in sEVs from CRC cells treated with 5-FU. Historically, the mechanism of action of 5-FU has been viewed as primarily affecting DNA synthesis due to misincorporation and inhibition of thymidine synthase (Wyatt and Wilson 2009). However, 5-FU also affects RNA metabolism leading to inhibition of processing of multiple classes of RNA and effects on ribosomes leading to altered translation efficiency of specific mRNAs and translational recoding (Mojardin et al. 2013; Ge et al. 2017; Chalabi-Dchar et al. 2021; Therizols et al. 2022). Consistent with effects on RNA metabolism, we find that treatment of cells with 5-FU leads to altered cellular and EV RNA profiles, including miRNAs, snRDs, and snoRDs. The data support that these effects are driven by 5-FU inhibition of pseudouridine modification because we detected significantly decreased levels of pseudouridine in purified EV RNA and because we observed enriched levels of uridine in RNAs that are differentially affected when comparing cells treated with 5-FU to untreated cells. Surprisingly, we did not observe significant differential effects on rDRs or tDRs, despite the fact that these RNAs are known to contain abundant levels of pseudouridine (Borchardt et al. 2020). Thus, the overall effects of 5-FU are pleiotropic and RNA-subclass dependent. Focusing on miRNAs, significant decreased detection in EVs may provide a potential biomarker for monitoring the effectiveness of 5-FU treatment during cancer chemotherapy. We chose to examine a heterogenous population of sEVs to not limit our analysis to specific EV classes and so that we could directly compare our data with previous RNAseq data from DLD-1 cells (Cha et al. 2015). For potential biomarker development, it may be preferable to analyze pseudouridine levels from a mix of EVs, but future work may define specific EV subclasses that are more affected by 5-FU treatment.

### Differential Effects of 5-FU on sRNA Expression and EV Export

Based on the abundance of pseudouridine modification, we expected to observe the greatest effect of 5-FU treatment on rRNA and tRNA. To our surprise, the effects of 5-FU exposure in DLD-1 cells were most dramatic for miRNAs, snDRs, and snoRDs. For miRNAs, 5-FU treatment led to a mix of both up- and down-regulated cellular expression levels of miRNAs when comparing treated to untreated cells. In contrast, there was a dramatic change in EV RNA profiles with significant inhibition of miRNA detection in EVs after 5-FU treatment. Remarkably, only 4 miRNAs were enriched in EVs after treatment with 5-FU, but 2 of those 4 are likely not bona fide miRNAs. The miRbase repository lists published miRNA sequences but does not curate or validate reported sequences (Kozomara and Griffiths-Jones 2011). To address this problem, the MirGeneDB applied additional criteria to better annotate bona fide miRNAs (Fromm et al. 2015) and concluded that 2 of the 4 up-regulated miRNAs in our differential EV expression analyses are not bona fide miRNAs. Only *miR-1249 and miR-576* remain as up-regulated in EVs after 5-FU treatment and these two had the lowest up-regulated fold-changes. Future work will be devoted to these miRNAs for why they are differentially affected by 5-FU.

For spliceosomal snRNAs, we also observed a decrease in EV snDR enrichment after 5-FU treatment. However, in contrast to cellular miRNA expression changes which were both up- and down-regulated, we observed only up-regulation of cellular snRNAs after 5-FU treatment. Based on that, one might predict increased detection of snDRs in EVs, but that is not what we observed. We found that most snDRs were down- regulated in EVs, suggesting a strong trafficking defect after 5-FU treatment

For snoRNAs, we observed a third pattern with both up-regulation of cellular expression and up-regulation of snoDRs in EVs after treatment with 5-FU. snoRNAs function to target rRNA for 2’-O-methylation and pseudouridine modification through RNA:RNA base pairing and the catalytic action of the DKC pseudouridine synthase (Huang et al. 2022). Given that we did not observe significant alteration in rDRs or tDRs in cells or EVs after 5-FU treatment, it seems likely that the increased expression and retention of snoDRs might be a cellular response to maintain pseudouridine levels in rRNA despite the presence of 5-FU.

### Pseudouridine Modification and Extracellular miRNA

The dramatic decrease in miRNA reads that we observed in EVs from cells treated with 5-FU raises several possible mechanisms of action for miRNA-mediated effects of 5-FU. The first is that pseudouridine modification may be an essential mark for EV export, perhaps in concert with specific-sequence motifs. This model predicts the presence of one or more RNA binding proteins that recognize pseudouridine residues (readers) and mediate miRNA export. To our knowledge, only 1 reader of pseudouridine has been proposed, a methionine aminoacyl tRNA synthetase (MetRS) that regulates translation (Levi and Arava 2021). Perhaps additional readers will be identified since pseudouridine is such a prominent modification. If not, an alternative role for pseudouridine modification in EV export could be related to RNA processing or RNA. It is known that pseudouridine modification can affect the structure, stability, and immunogenicity of RNA (reviewed in (Borchardt et al. 2020)) and was critical to the success of the COVID-19 mRNA vaccines (Morais et al. 2021). Previous reports have shown that pseudouridine formation can regulate miRNA processing and also that incorporation of 5-FU can inhibit U2 snRNA modification and assembly (Patton 1991; Kurimoto et al. 2020; Song et al. 2020). Thus, incorporation of 5-FU could block pseudouridine modification of miRNAs and snRNAs leading to decreased processing and intracellular accumulation of precursor RNAs. For miRNAs, incomplete processing might lead to accumulation of more stable double stranded RNAs which could then also lead to intracellular accumulation.

### Sensitivity and Resistance to 5-FU

One explanation for why we did not observe down-regulation of rDRs and tDRs could be due to differential inhibition of pseudouridine synthases by 5-FU. There are 13 pseudouridine synthases in human cells (Jin et al. 2022) and not all of them are inhibited by 5-FU incorporation, indicating that these synthases are not necessarily functionally redundant (Spedaliere and Mueller 2004). The synthases that act on rRNA or tRNA might not overlap with those that act on miRNAs and snRNAs. Also, it is important to recognize that there will be differences in expression and localization depending on the concentration and duration of 5-FU exposure. We subjected DLD-1 cells to a relatively moderate amount of 5-FU for a limited time to observe initial changes in small RNA expression and localization. Future experiments are planned to compare transient and long-term exposure to 5-FU.

Consistent with differential activity among pseudouridine synthases, chemotherapeutic resistance to 5-FU is often associated with increased expression of one of more pseudouridine synthases (Jin et al. 2022). In hepatocellular carcinoma and colorectal cancer, higher expression of pseudouridine synthases (PUS1, PUS7, PusL1, RPUSD3, DKC1, PUS7L, PUS10, and RPUSD1) correlates with poor prognosis (Jin et al. 2022; Liang et al. 2022; Zhang et al. 2023). Likewise, overexpression of the Dyskerin pseudouridine synthase that acts on rRNA (DKC1) predicts poor prognosis in breast cancer, while knockdown of DKC1 and Mek1/2 restricts colorectal cancer cell growth (Elsharawy et al. 2020; Kan et al. 2021). These studies are also consistent with work from *S. cerevisiae* where 5-FU toxicity requires Cbf5p, a pseudouridine synthase (Hoskins and Butler 2008). Data mining of published RNAseq data sets from 5-FU resistant cells showed that 6 pseudouridine synthases were up-regulated (PUS3, PUS7, PUS7L, Trub1, RPUSD3, and RPUSD1) with the remaining 7 being either unaffected or down-regulated (Chauvin et al. 2022). Analysis of a 5-FU resistant colorectal cancer line (HCT116) found a negative correlation between 5-FU sensitivity and PUS1 levels and stabilization of PUS1, PUS7, PUS10, and TRUB1 in mouse xenografts (Liang et al. 2022). In our RNAseq data after short term exposure of DLD-1 to 10μM 5-FU, we found that PUS7, TRUB1, TRUB2, RPUSD1, PUS10, and DKC1 were up-regulated with the other pseudouridine synthases either unaffected (PUS1, PUS 7L, PUSL1) or down- regulated (PUS3, RPUSD3, RPUSD4, and RPUSD2). Together, the data indicate that not all pseudouridine synthases are equivalent and raise the possibility that upon exposure to 5-FU, cells adapt expression levels among the 13 members of the family to survive.

### RNA Base Modifications and EV Export

PTxMs on cellular and extracellular sRNAs provide another layer of gene regulation. 5-FU treatment significantly decreased the levels of pseudouridine in both cells and EVs, as quantified using LC-MS/MS. While we focused on pseudouridine because of inhibition by 5-FU, we also detected changes in base modifications of adenosine, guanosine, cytosine, and inosine with significant increases in post-transcriptional methylation on cellular sRNAs including m^1^A, m^1^I, m^3^C, m^5^C, m^1^G, m^2^G, m^7^G, and m_2_ ^2^G. This is the first analysis of PTxMs using mass spectrometry on both cellular and secreted EV sRNAs and will be an important powerful tool for future studies on sRNA trafficking and localization. Because pseudouridine was the only PTxM observed to be altered in sEVs after 5-FU treatment, the LC-MS/MS data support a role for pseudouridine in sRNA selection and/or export to EVs.

## Acknowledgements

The authors would like to thank all members of the Coffey, Weaver, and Patton labs for advice and suggestions.

## Author Contributions

SQ, HN, ES, XL, DM, and CM generated all data in the manuscript. SQ, HN, QL, JLF, JK, AMW, and RJC provided advice and suggestions. KJC and JGP wrote the paper with helpful edits from AMW, SQ, and HN.

## Figure Legends and Tables

Table S1. Long RNAseq data from DLD-1 cells exposed to 5-FU. Sequencing data for libraries generated from RNA >200nt isolated from cells exposed to 5-FU (FiveFU) or not (SF). Significant differentially expressed genes with absolute fold change > 2 or < 0.5, and adjusted p values (padj) <0.05 were determined using DESeq2 (v1.30.1)(Love et al. 2014).

Table S2. Differentially expressed genes in cells exposed to 5-FU. Table showing the identity of significant differentially expressed genes with absolute fold change > 2 or < 0.5, and adjusted p values (padj) <0.05 when comparing treated to untreated DLD-1 cells. Differential expression was determined using DESeq2 (v1.30.1)(Love et al. 2014).

Table S3. Small RNAseq data from DLD-1 cells and EVs exposed to 5-FU. Sequencing data for libraries generated from RNA <200nt isolated from cells and EVs exposed to 5-FU (FiveFU) or not (SF). Significant differentially expressed genes with absolute fold change > 2 or < 0.5, and adjusted p values (padj) <0.05 were determined using DESeq2 (v1.30.1)(Love et al. 2014). Tabs show sequencing data for miRNAs, rDRs, tDRs, snDRs, snoDRs and yDRs from both cells and EVs.

Table S4. 5-FU treatment primarily reduces EV miRNA expression. After 5-FU treatment, the majority (278) of miRNAs showed significant changes in expression in EVs (blue) with no significant change in cellular expression (red). 13 miRNAs showed significant changes in cellular expression (red) but not in EVs (blue). 15 miRNAs showed significant changes in both cellular (red) and EV (blue) expression after 5-FU treatment.

---

**Figure S1.**
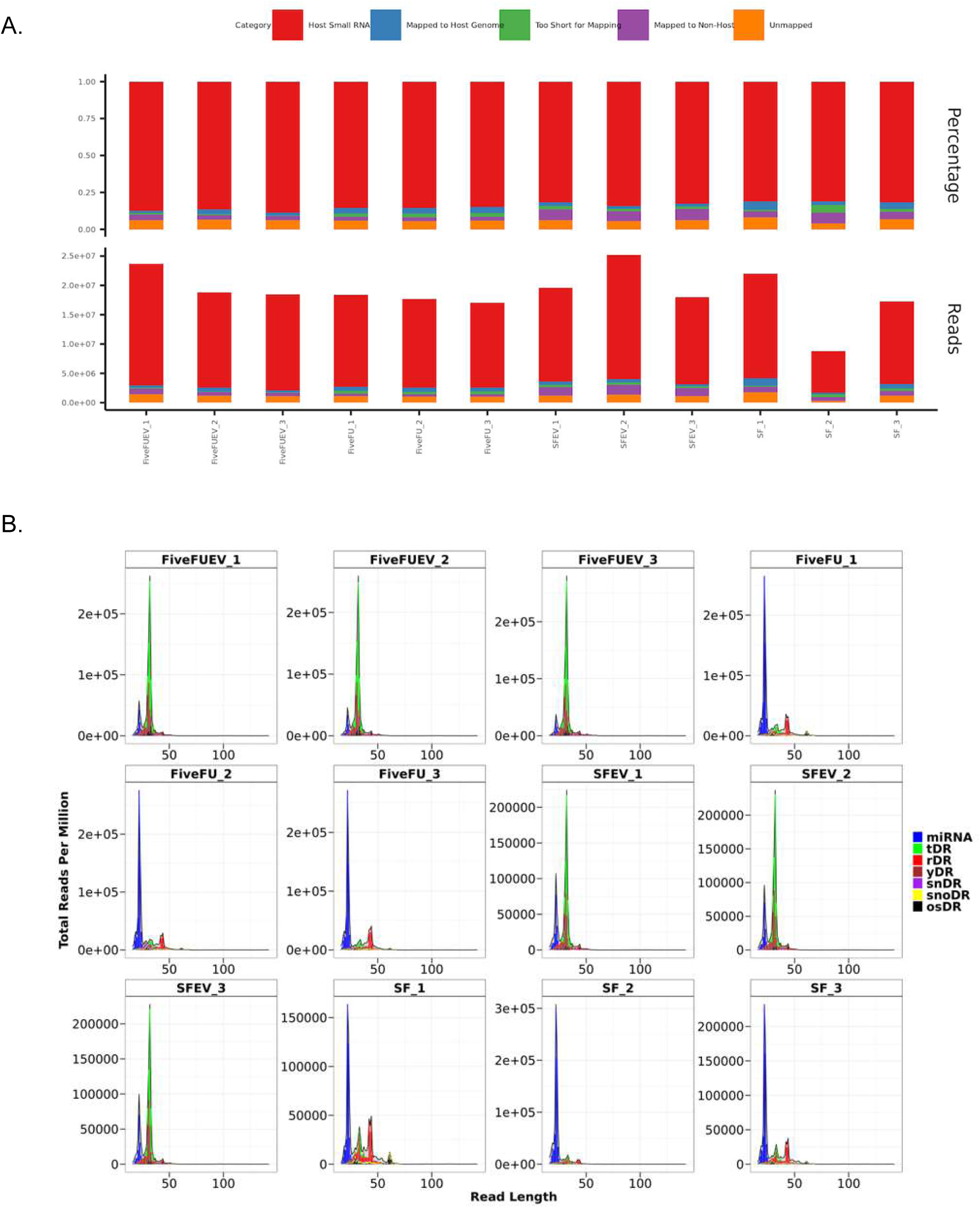
Quality control of small RNAseq. RNAseq was performed in triplicate on RNA (<200nt) isolated from DLD-1 cells and EVs treated (FiveFU) or not (SF) with 5-FU. **A.** Read numbers and percentage of reads mapping to the categories indicated at the top are as shown. **B.** Read size distribution from the same biological replicates in A with the indicated RNA subclass including miRNA, tRNA-derived fragments (tDR), rRNA-derived fragments (rDR), Y RNA-derived fragments (y DRs), spliceosomal snRNA-derived fragments (snDR), snoRNA-derived fragments (snoDR), and other assorted miscellaneous small RNA fragments (osDR).

**Figure S2.**
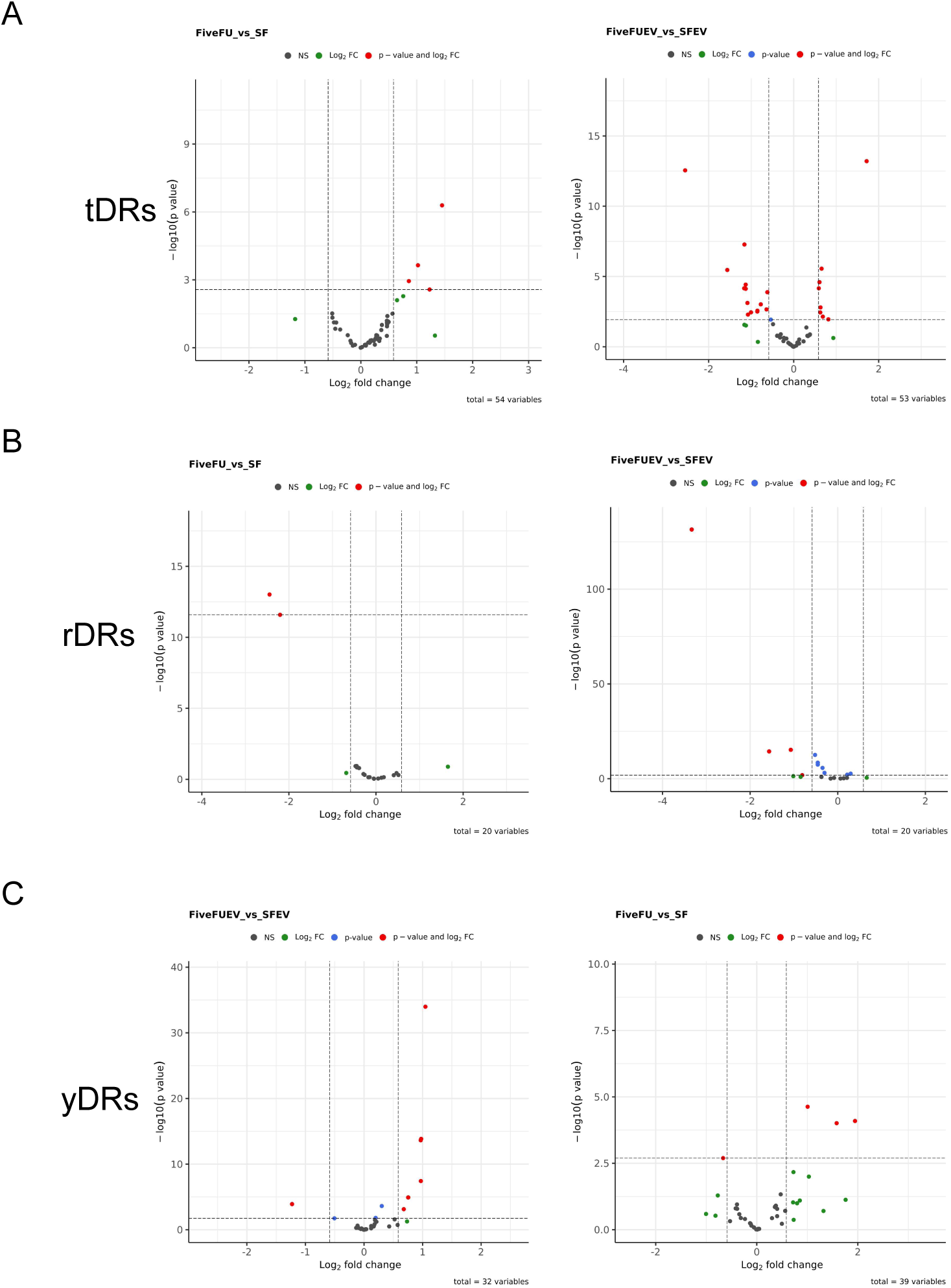
Differentially expressed tDRs and rDRs comparing DLD-1 cells and EVs grown in the presence or absence of 5-FU. Volcano plots showing up- and down-regulated tDRs (**A**), rDRs (**B**), and yDRs (**C**) in cells and EVs treated (FiveFU) or not (SF) with 5-FU. Individual data points represent transcripts with fold-changes and significance as in Fig. 1.

**Figure S3.**
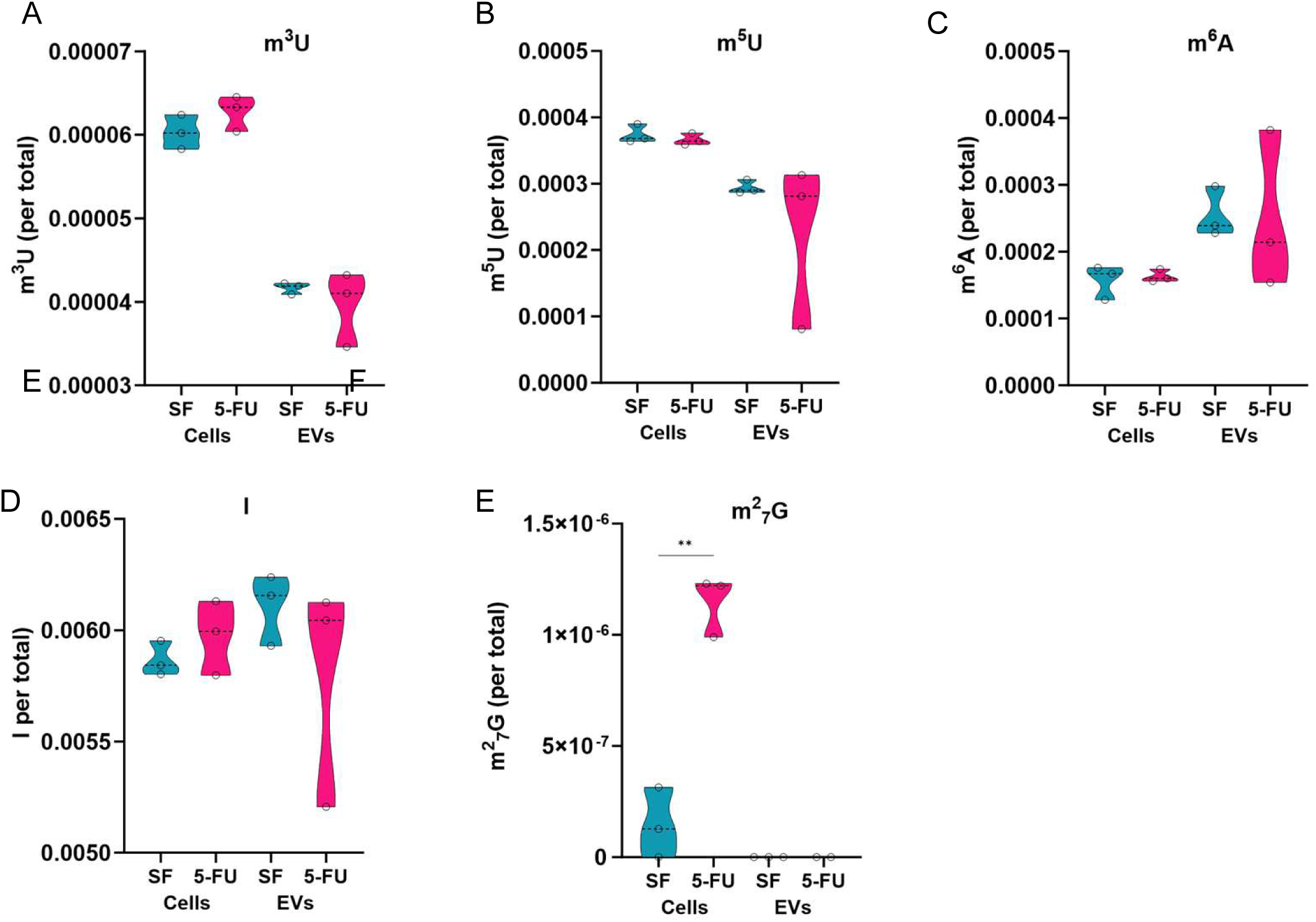
Effects of 5-FU on RNA base modification in cells and EVs. LC-MS/MS approaches were used to quantify RNA base modifications in RNA from cells and secreted sEVs. **A-E**. Quantitation of the effect of the indicated base modifications in cells and EVs treated or not with 5-FU. Base modifications include methylated uridine (m^3^U, m^5^U), methylated adenosine (m^6^A), Inosine, and dimethylated guanosine (m^2^_7_G).

